# Rapid Selectivity to Natural Images Across Layers of Primate V1

**DOI:** 10.1101/2022.01.23.477422

**Authors:** Xiaomo Chen, Shude Zhu, Kunlun Bai, Ruobing Xia, Nathan C.L. Kong, Anthony M Norcia, Tirin Moore

## Abstract

Visual systems are thought to have adapted to the statistical properties of natural scenes. However, the extent to which visual neurons respond selectively to natural images, and the stage at which that selectivity emerges remains unclear. To address these questions, we recorded the visual activity of neurons in macaque V1 using high-density electrode arrays (Neuropixels), and compared neuronal responses to images presented at three levels of naturalness. We found that within 60 ms of stimulus onset, neurons in all cortical layers, including input layers 4C alpha and beta, responded more vigorously to natural images than to statistically matched naturalistic texture and noise images. The result remained when residual variations in the local image statistics were factored out. V1 neurons also showed high population and lifetime sparseness for natural images. Across the population of V1 neurons, sensitivity to natural images exceeded the sensitivity to other image categories. The results reveal a rapid and pervasive preference for natural images is present at the earliest stages of cortical processing.

## Introduction

Neurons in the visual system are believed to have adapted to the statistical properties of the natural environment ^1,2^. Along the visual hierarchy, neurons in the primary visual cortex (V1) represent information about local edge elements such as their local orientation and spatial scale ^3–6^. This representation of local edge elements has long been thought to be subsequently combined to construct corners, junctions, and more extensive contours, eventually leading to shapes, forms, and objects in later stages along the ventral stream ^7–10^. Along this presumptive visual hierarchy, it is still unclear at which processing stage visual neurons become sensitive to the structure of natural images.

A fruitful approach to studying natural images processing has been to compare responses to synthetic images matched on various summary statistics to those of natural images ^11–14^. One of these synthetic image types comprises images that match the power spectrum of corresponding natural images, spectrum-matched noise images (SMNIs). These images preserve a low-level summary statistic of natural images – the power spectrum. A second type of synthetic image, naturalistic texture images (NTIs), additionally maintains several pair-wise correlations among the outputs of V1-like filters ^15^, capturing mid-level structure present in natural images. This latter approach was first used to study neuronal responses at the late stages of visual processing in macaque cortex where it was found that single neurons in V4 and IT show higher sensitivity to natural images than naturalistic textures ^12,13,16^. Later work in V1 and V2 found that single V1 neurons are insensitive to the pair-wise correlation structure present in NTIs, and that sensitivity to this mid-level structure first emerges in V2 ^11,14,17^. A study in human visual cortex compared BOLD responses to SMNIs, NTIs, and natural images and found an orderly progression where selectivity to natural images emerged first in areas anterior to V3, such as V4, while selectivity to natural textures emerged in V2 ^17,18^, consistent with the single-neuron data in macaque. A parsimonious conclusion from the human study above is that selectivity to natural images is not present in V1.

However, other studies using natural images have shown that V1 neurons exhibit high selectivity and response sparseness for natural images in primates, rodents and ferrets ^19–23^, including when compared to SMNIs ^19–21^. As different image sets, species and response metrics (e.g., mean firing rates vs spike count correlations) have been used in prior studies, the compatibility of these two lines of research remains unclear, and the selectivity of V1 neurons to natural images thus remains unresolved. Here we recorded the responses of approximately 622 V1 neurons across the layers using high-density electrode arrays that were evoked by a large set of images containing three levels of naturalness: 300 natural images, 300 NTIs, and 300 SMNIs (Methods). We observed that V1 neurons responded more vigorously to natural images than to naturalistic texture and noise images within 60ms of stimulus onset. The result remained when variations in local image statistics were controlled by generalized linear regression models. This sensitivity to naturalistic images was robustly observed throughout all cortical layers, including neurons in input granular layer 4C alpha and beta. Moreover, on the population level, V1 neurons showed high population and lifetime sparseness, and were less synchronized with the overall population activity when responding to natural images, suggesting a more efficient coding strategy. Furthermore, neuronal ensembles exhibited higher discriminability for natural images. The results reveal a rapid and pervasive preference for natural images is present at the earliest processing stages of visual cortex.

## Results

We recorded the activity of single neurons in area V1 in 2 anesthetized monkeys using high-density, multi-contact Neuropixel probes (version 3A; IMEC Inc, Belgium) (Fig. 1a and b). In total, the activity of 622 isolated single neurons was recorded across the layers of V1 in 4 penetrations.

**Figure 1.**
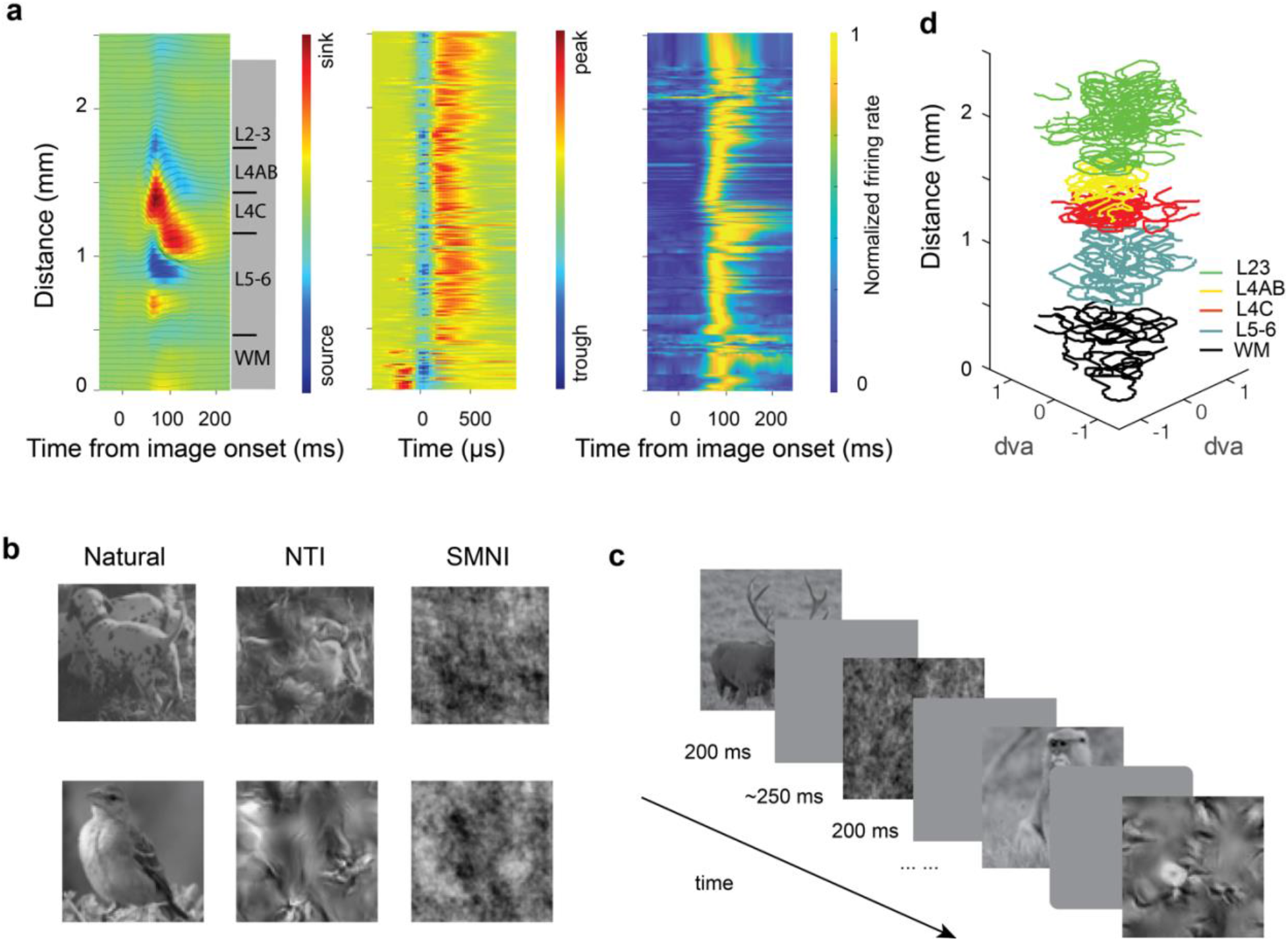
Experimental setup and an example session. (**a**) An example recording session. Current source density analysis was used to identify cortical layers (left); thin lines indicate the borders of different cortical layers. Spike waveform voltage traces across cortical depth (middle). Average neuronal responses simultaneously recorded across layers to all images in one experiment session (right). **(b**) Visual receptive fields (RFs) reconstructed across layers in one example recording session. Different colors indicate RFs reconstructed in the identified laminar compartments. (**c**) Illustration of the stimulus presentation. Each image was presented for 200 ms followed by a 250-ms gray background. (**d**) Example natural images (left), naturalistic texture images (middle), and spectrum-matched noise images (right).

For each recording, we estimated the borders of laminar compartments by combining the histological data with current-source density (CSD) measurements (Methods) (Fig. 1a). To assess the sensitivity of V1 neurons to image naturalness, we compared neuronal responses to three sets of images that varied in their higher-order statistics, specifically natural images, naturalistic texture images (NTIs), and spectrum-matched noise images (SMNIs) (Methods) (Fig. 1c and d and Supplementary Fig. 1).

### V1 neurons respond more vigorously to natural images

We first compared the magnitude of V1 responses to different levels of image naturalness. We found that V1 neurons responded more vigorously to natural images compared to synthetic images (Fig. 2a). First, we measured the entire neuronal response (40 - 200 ms from image onset) and calculated a modulation index to quantify response differences between different image types (Methods). Consistent with previous studies ^11,14^, we found no significant difference between the magnitude of responses to NTIs and to SMNIs (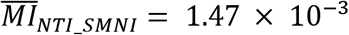, *p* = 0.67, paired t-test). However, the neuronal responses to natural images were consistently larger than those to NTIs (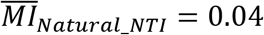, *p* < 10^−31^, t-test) and to SMNIs (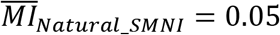, *p* < 10^−42^, paired t-test). The differences in response to natural images amounted to an ~11% increase above responses to synthetic images. This effect was reliably observed in both monkeys (Supplementary Fig. 2). Similar results were observed when we further divided neuronal responses into early (40 – 100ms) and late (100 – 200ms) response epochs. There was no significant difference between the magnitude of responses to the two synthetic images in either the early (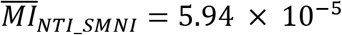, *p* = 0.99, paired t-test) or late epochs (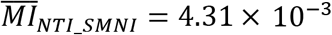, *p* = 0.31, paired t-test). In contrast, natural image responses were greater than those of synthetic images in both the early (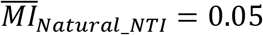, *p* < 10^−26^; 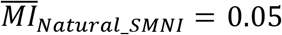, *p* < 10^−30^, paired t-test) and late epochs (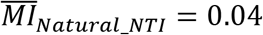, *p* < 10^−19^; 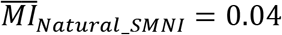, *p* < 10^−30^, paired t-test) (Fig. 2b and c). We further examined the variance of neuronal responses to different levels of naturalness by calculating modulation indices for the Fano factor of neuronal responses. In contrast to the firing rate, we observed no significant differences in Fano factor between the image conditions (Fig. 2c). Thus, natural images elicited larger responses throughout early and late epochs, but responses exhibited similar variability.

**Figure 2.**
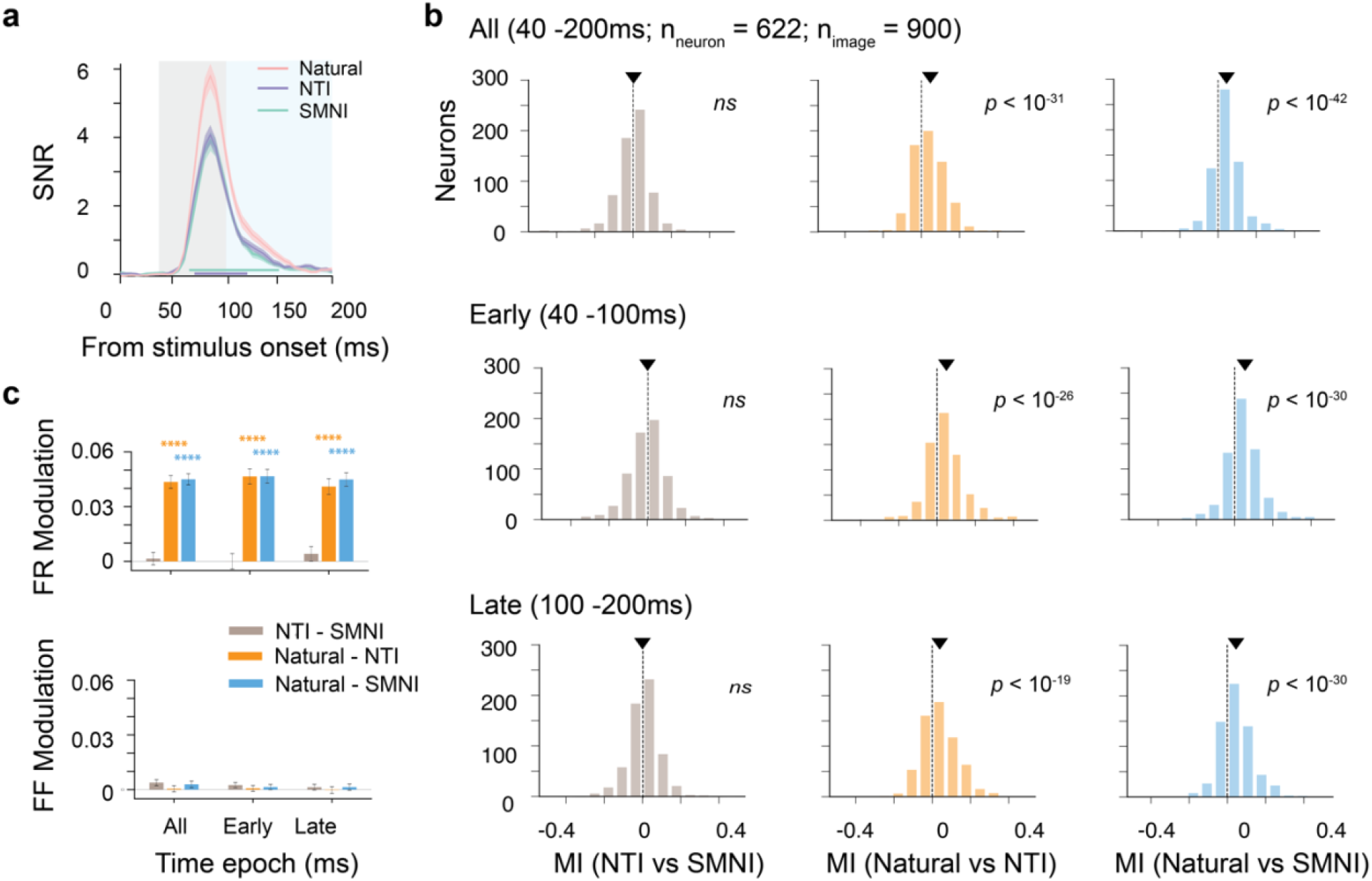
Neurons respond more vigorously to natural images than to synthetic images. (**a**) Mean responses of all simultaneously recorded neurons to different image categories: natural images (pink), NTIs (purple), and SMNIs (green). The thickness of lines indicates s.e.m.. across images within one category. Thick lines at the bottom indicate the time period when responses significantly differ (natural images vs NTIs: purple; natural images vs SMNIs: tiffany). The shaded areas indicate the early (gray), late (blue) and all (gray and blue) time epochs. (**b**) Histograms of modulation indices. The modulation index was computed separately for comparison between NTIs and SMNIs (left), natural images and NTIs (middle), and natural images and SMNI (right). (**c**) Bar plots summarize the firing rate modulation index (FR Modulation) and Fano factor modulation index (FF Modulation) during different response epochs. Asterisks denote significant differences between different conditions (\*\*\*\**p* < 10^−4^).

### A generalized linear model (GLM) control for local contrasts

Although V1 neurons appeared to be selective for natural images, residual low-level factors might have contributed to the observed response differences. Specifically, we considered that differences in local RMS contrast and band-limited contrast (Fig. 3a) could potentially be a source of the enhanced responses to natural images. Although the three image sets are identical in several global image statistics, small differences in RMS contrast between natural and synthetic images (Fig. 3b) and in band-limited contrast between natural and the NTIs (Fig. 3c) may nonetheless persist.

**Figure 3.**
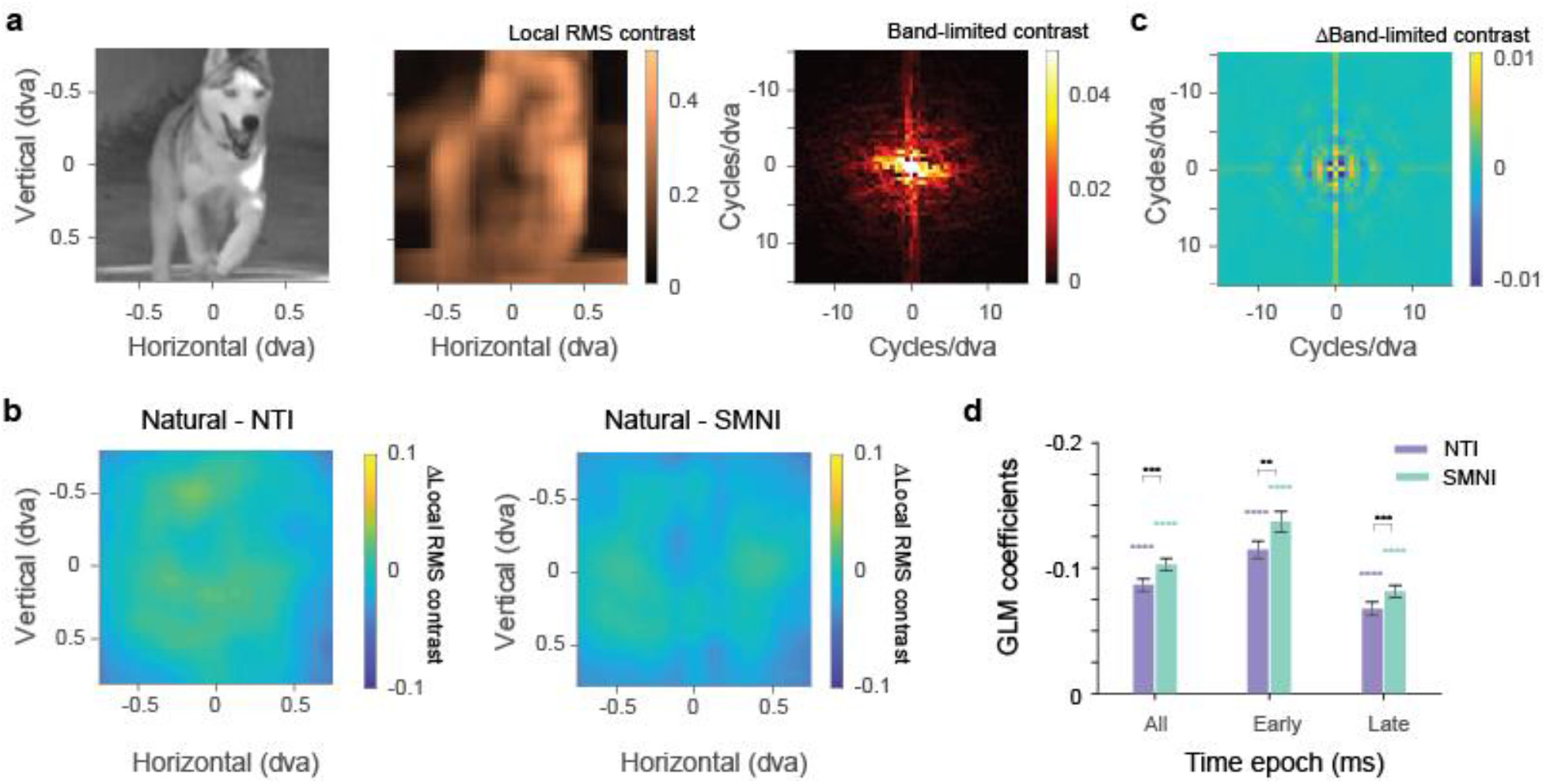
Controlling the effects of local contrasts. (**a**) An example natural image, its local RMS contrast (middle) and band-limited contrast (right). (**b**) Differences of average RMS contrast between natural images and NTIs (left), and between natural and SMNIs (right). (**c**) Differences of average band-limited contrast between natural images and NTIs. (**d**) Bar plot summarizes the GLM coefficients for comparisons to NTIs and SMNIs during different time epochs. Asterisks denote significant differences between the natural image condition and the NTI and SMNI conditions (\**p* < 0.05, \*\**p* < 10^−2^, \*\*\**p* < 10^−3^, and \*\*\*\**p* < 10^−4^). Error bars denote the s.e.m..

To control for the contribution of these residual image statistic differences, we developed a GLM that modeled the activity of individual neurons based on the variables that could affect activity, namely band-limited contrast, local RMS contrast, and image category (Methods). When RMS and band-limited contrasts were controlled for, we found that responses to natural images still differed from those to synthetic ones. GLM coefficients for image categories were significantly negative for both SMTIs and NTIs (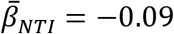, *p* < 10^−59^; 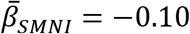, *p* < 10^−75^, t-test) indicating larger responses to natural images. In addition, there were small, but significant, differences between the NTI and SMNI coefficients (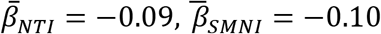, *p* < 10^−3^, paired t-test) indicating that the preference for natural images was larger for the SMNIs. Similar results were observed for both early (40 – 100ms) (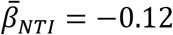, *p* < 10^−51^; 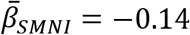, *p* < 10^−53^, t-test) and late response epochs (100 – 200ms) (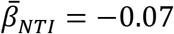, *p* < 10^−30^; 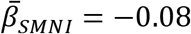, *p* < 10^−49^, t-test) (Fig. 3d and c). Moreover, the early responses exhibited a greater preference for natural images than the later ones (NTI: 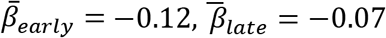, *p* < 10^−15^; SMNI: 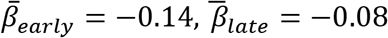, *p* < 10^−12^, paired t-test) suggesting that the natural image selectivity is present at the earliest stages of processing within V1.

### Selectivity to natural images throughout cortical layers

To further explore the possible origins of the preference for natural images, we next examined the spatiotemporal pattern of sensitivity to natural images across different cortical layers. We considered that if the sensitivity to naturalness in V1 originates from feedback from later visual cortical stages, e.g., V2, then we should observe stronger and earlier developing sensitivity within supragranular and infragranular layers, with weaker and slower developing sensitivity within granular layer 4C ^14,24,25^. Using the simultaneously recorded activity, we compared the sensitivity to natural images for neurons distributed across laminar compartments (Fig. 4a). We found that selectivity to natural images was present across all layer compartments in V1. Neurons in all cortical layers, including input layer 4C, responded more vigorously to natural images than to synthetic ones. We calculated the GLM coefficients for neurons in each laminar compartment in both early and late epochs (Fig. 4b). GLM coefficients revealed a significant preference for natural images across all layers when compared to NTIs in both early and late epochs (early: 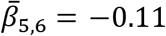, *p* < 10^−8^; 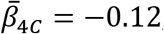, *p* < 10^−39^; 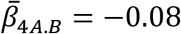, *p* < 10^−8^; 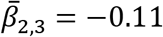, *p* < 10^−16^; late: 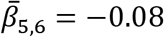, *p* < 10^−4^; 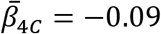, *p* < 10^−22^; 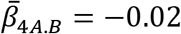, *p* < 0.05; 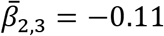, *p* < 10^−12^, t-test), and when compared to SMNIs (early: 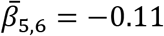, *p* < 10^−9^; 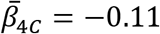, *p* < 10^−37^; 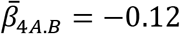, *p* < 10^−12^; 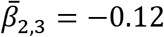, *p* < 10^−19^; late: 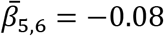, *p* < 10^−5^; 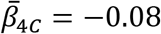, *p* < 10^−22^; 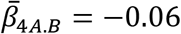, *p* < 10^−8^; 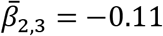, *p* < 10^−13^, t-test). In addition, neurons recorded within the white matter showed a similar effect (early: 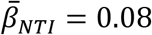, *p* < 10^−4^; 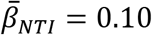, *p* < 10^−5^; late: 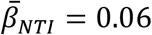, *p* < 10^−3^; 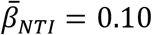, *p* < 10^−5^, t-test). To examine the time course of sensitivity across layers, we calculated GLM coefficients across the full response time window (Methods). These coefficients revealed significant sensitivity within ~60 ms of stimulus onset across layers for both NTIs and SMNI (Fig. 4c). Moreover, the largest proportion of neurons with an earlier onset times resided in Layers 4A/B and 4C, a difference that was greater than that of Layer 2-3 for both NTIs (Layer 4A/B: *p* = 0.08; Layer 4C: *p* = 0.03, K-S test) and SMNIs (Layer 4A/B: *p* < 10^−5^; Layer 4C: *p* < 10^−3^, K-S test) (Fig. 4d). We further subdivided Layer 4C into L4C*α* and L4C*β* to examine the difference between the 2 input sublaminae (Methods) and found that a similar pattern of similar pattern of sensitivity was observed in both L4C*α* and L4C*β* (Supplementary Fig. 3). Thus, the preference for natural images emerged early in V1 and to a greater extent within input layers.

**Figure 4.**
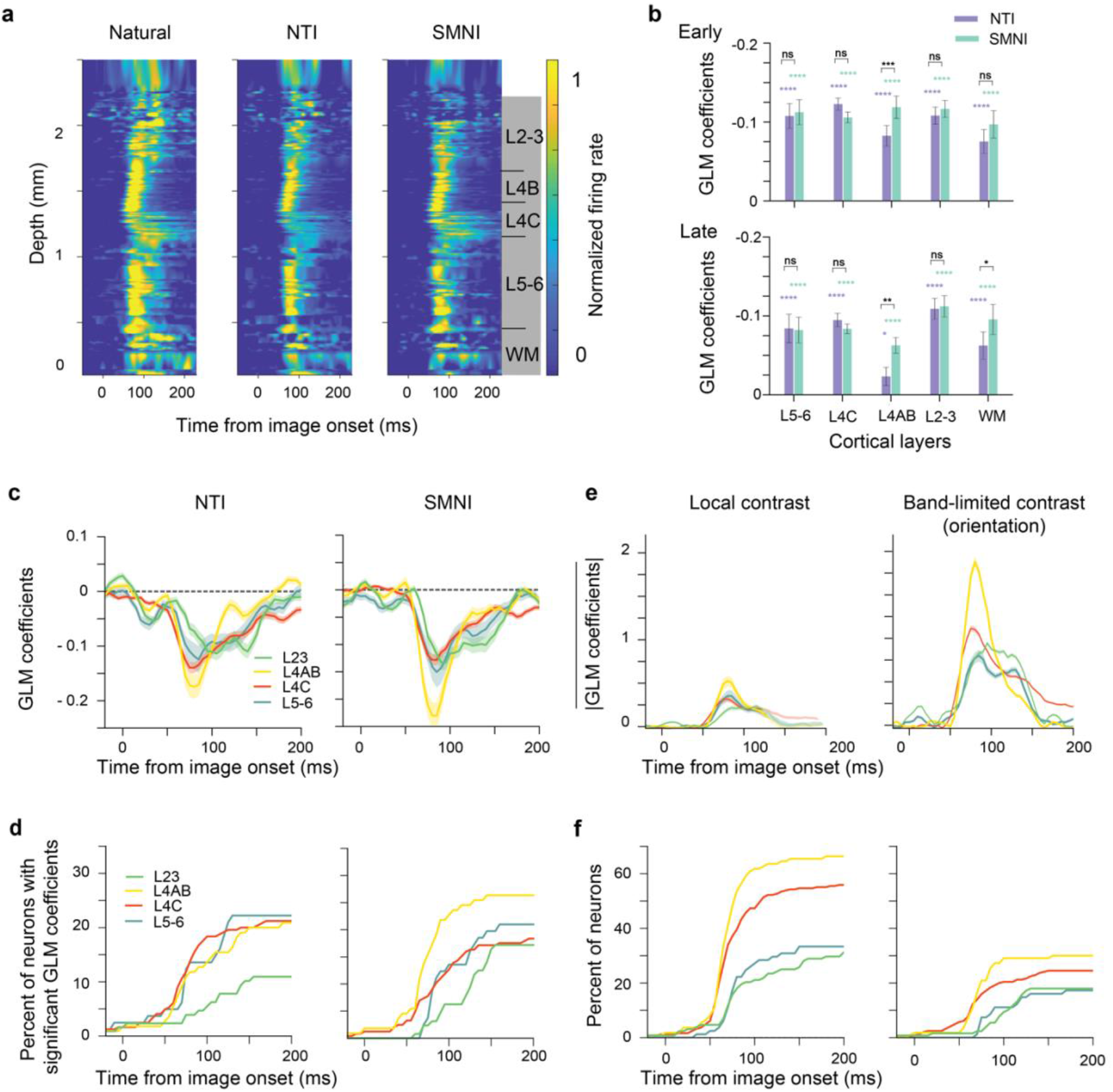
Sensitivity to natural images across cortical layers. (**a**) Average neuronal responses simultaneously recorded across layers for Natural images (left), NTIs (middle), and SMNIs (right) in a representative recording session. (**b**) Bar plots summarize the GLM coefficients for comparisons of natural image responses to NTIs and SMNIs across cortical layers. (**c**) Average GLM coefficients for comparisons to NTI images (left) and SMNI images (right) as a function of time, separated by different cortical layers. Shading around the lines indicates the s.e.m. of GLM coefficients across neurons. (**d**) Percentage of Neurons with significant GLM coefficients for comparisons to NTI images (left) and SMNI images (right), separated by different cortical layers. (**e**) Average GLM coefficients for local contrast (left) and orientation (right) as a function of time, separated by different cortical layers. (**f**) Percentage of Neurons with significant GLM coefficients for local contrast (left) and orientation (right).

In addition, we examined the time course of GLM coefficients for local contrast modulation and band-limited contrast. As we discretized the band-limited contrast into sectors based on orientations (Methods), sensitivity to band-limited contrast represents orientation sensitivity. Given that we expect that contrast and orientation sensitivity in V1 are not driven by a feedback modulation from higher visual areas, we could compare the time course of contrast and orientation sensitivity to that of sensitivity to image naturalness. Both local contrast and orientation sensitivity showed similar temporal profiles (Fig. 4e and f). Specifically, the coefficients revealed significant sensitivity within 50-60 ms of stimulus onset across layers for both contrast and orientation. In addition, Layer 4A/B showed an earlier sensitivity to local contrast and orientation than both Layer 2-3 (local contrast: *p* < 10^−2^; orientation: *p* < 10^−4^, K-S test) and Layer 5-6 (local contrast: *p* = 0.05; orientation: *p* = 0.04, K-S test). Moreover, the input Layer 4C also showed an earlier sensitivity to local contrast and orientation than Layer 2-3 (local contrast: *p* < 10^−2^; orientation: *p* < 10^−2^, K-S test) and but not Layer 5-6 (local contrast: *p* = 0.07; orientation: *p* = 0.06, K-S test). Thus, as with contrast and orientation, natural image selectivity emerges earliest within the input layers of V1.

### Sparse coding of natural images

After quantifying natural image selectivity for individual neurons in V1, we examined the response to forms of response sparseness: lifetime sparseness and population sparseness ^26^. Lifetime sparseness quantifies how sparsely a single neuron responds to different images ^27^. Consistent with previous studies in cats ^28,29^, rodents ^21,30^, ferrets ^31,32^ and primates ^22,23,33^, we observed that single neurons in V1 respond more selectively to natural images compared to synthetic images. For a given number of images, significantly more response variance is explained by natural images than by synthetic images (Fig. 5a and Supplementary Fig. 4a). Fewer images are needed to capture 80% of the response variance with natural images than with synthetic images (*p_natural vs NTIs_* < 10^−95^; *p_natural vs SPNIs_* < 10^−107^, paired t-test) (Methods). This result indicates that V1 neurons tend to respond individually to a more specific set of natural images than they do to synthetic images.

**Figure 5.**
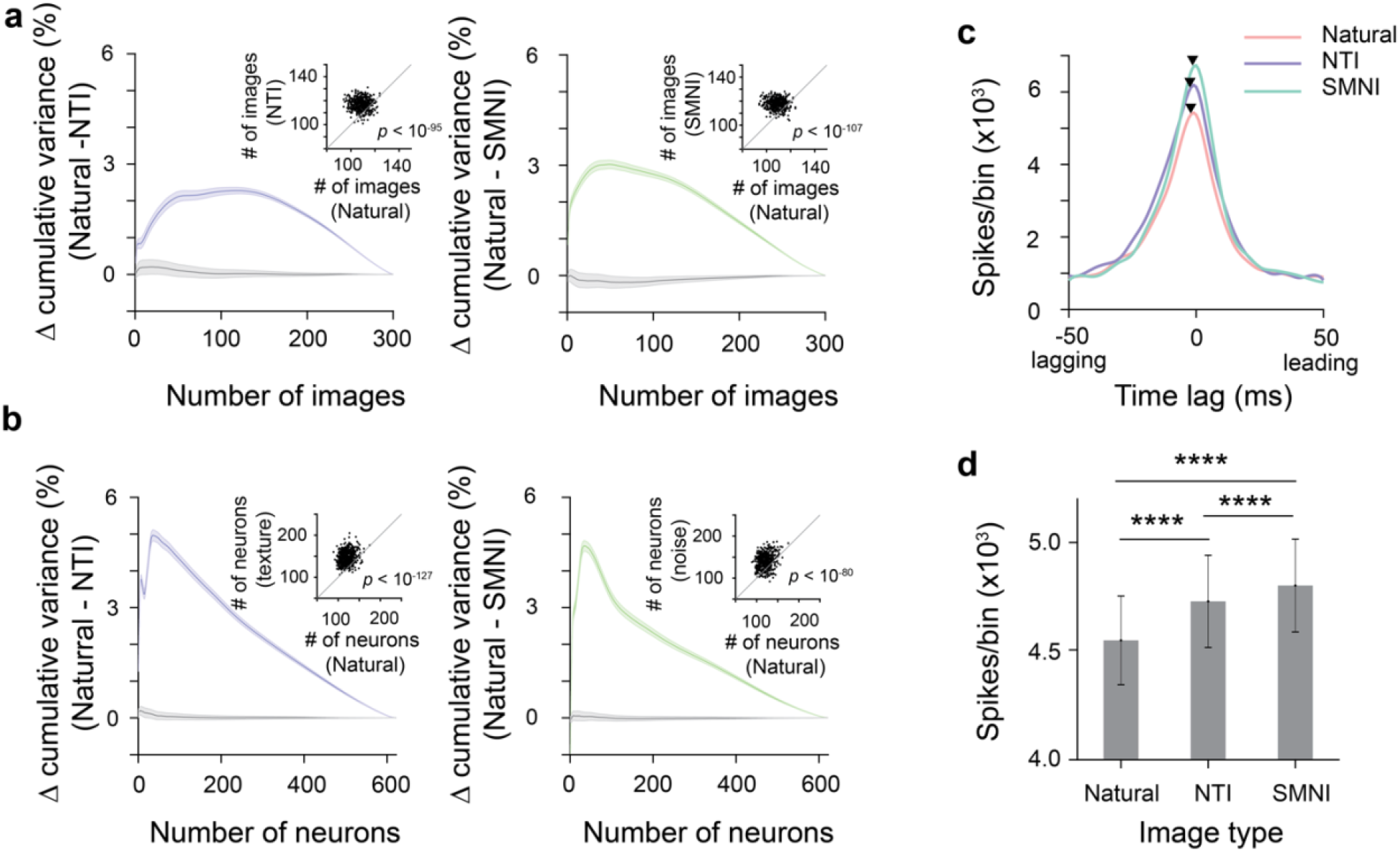
Sparseness and population synchrony. (**a**) Lifetime sparseness: Differences in cumulative variance explained by different numbers of images. Scatter plots compare the number of images needed to capture 80% of the response variance for natural images and synthetic images. Left, comparison between natural images and NTI. Right, comparison between natural images and SMNI. The gray lines represent the difference in cumulative variance when the labels for different image sets were shuffled (N = 1000). The thickness of lines indicates s.e.m.. (**b**) Population sparseness: Differences in cumulative variance explained by different numbers of neurons. Scatter plots compare the number of neurons needed to capture 80% of the response variance for natural images and synthetic images. Left, comparison between natural images and NTI. Same conventions as (a). (**c**). A representative spike time cross-correlation between a single neuron and the population response for natural images (Nature, red), synthetic texture images (NTI, purple), and spectrally matched noise images (SMNI, green). (**d**). The bar graph shows the average peak magnitude of cross-correlogram for different image categories. Asterisks denote significant differences between different conditions (****, *p* < 10^−4^). Error bars denote the s.e.m.

Next, we examined the population sensitivity to different levels of naturalness. Previous studies in cats ^29^, rodents ^21,30^, ferrets ^32^, and primates ^22^ have shown that, within a population, only a small subset of the V1 neurons within the recorded actively encode a given natural image. These results shows that V1 neurons respond sparsely to natural images and thus show population sparseness. In this study, we quantified the population sparseness of the V1 neurons in our high-density recordings by computing the cumulative variance as a function of the number of neurons. We observed that, for a given number of neurons, significantly more response variance is explained with natural images than with synthetic images (Fig. 5b and Supplementary Fig. 4a). Consequently, we observed that fewer neurons are needed to capture 80% of the total variance of the recorded neurons (*p_natural vs NTIs_* < 10^−127^; *p_natural vs SPNIs_* < 10^−80^, paired t-test) (Methods) (Fig. 5b and Supplementary Fig. 4b). This result suggests that fewer neurons are driven by a given stimulus in the natural image set than a stimulus in the synthetic image sets.

We also asked whether the population sparseness of natural image responses extends to a finer temporal domain. Specifically, we tested whether different image sets exhibited different patterns of population synchrony. To measure population synchrony, we computed the ‘coupling’, the spike time correlation between individual V1 neurons and the simultaneously measured population activity within the column ^34^. To control for differences in the magnitude of responses to different image categories, we mean-matched the neuronal response across image conditions. We further used a jittered correlation method to compute spike-count correlations between individual neuronal responses and population responses (Methods). Although the time lag of population coupling was similar across image types (Supplementary Fig. 5a), we found that the strength of the population coupling was inversely related to the level of naturalness (Fig. 5c and Supplementary Fig. 5b). Thus, in this finer temporal domain, our result shows that individual V1 neurons were less synchronized with the local neuronal population when responding to natural images, (Fig. 5d) (Natual vs NTI: *p* < 10^−22^; Natural vs SMNI: *p* < 10^−34^; NTI vs SMNI: *p* < 10^−4^, paired t-test).

### Robust population representation of natural images

Next, we tested whether the differential responses to image naturalness resulted in greater discriminability among natural images. We computed discriminability indices (*d*′) from the evoked responses of all the neurons to all image pairs within each image set ^35,36^. Discriminability is related to the Fisher information that the population ensemble provides about stimulus identity ^35–37^. To enable us to determine *d*′ accurately and reduce the noise of the measurement, we performed a dimensionality reduction by using PCA on the averaged firing rates for each image, within image category. We then identified and retained only ten population vector dimensions in which the image-responses were highly distinguishable. This analysis was done separately for neuronal responses to the different image categories. For each image category, *d*′ was sorted so that the largest *d*′was plotted in the top-left quadrant and the smallest *d*′was plotted in the bottom-right quadrant (Fig. 6a). In this reduced dimension representation, the population code for natural images had the largest mean *d*′ 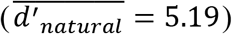. Therefore, the representations of natural images had the highest discriminability between all image pairs compared to those of synthetic images (Fig. 6b). Like the results observed in the first ten PCs, the population code for natural images has the largest mean *d*′ across all first ten PC dimensions (Supplementary Fig. 6). In addition, there was a small but significant difference between NTIs and SMNIs (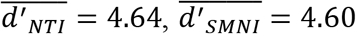, p < 10^−2^, paired t-test).

**Figure 6.**
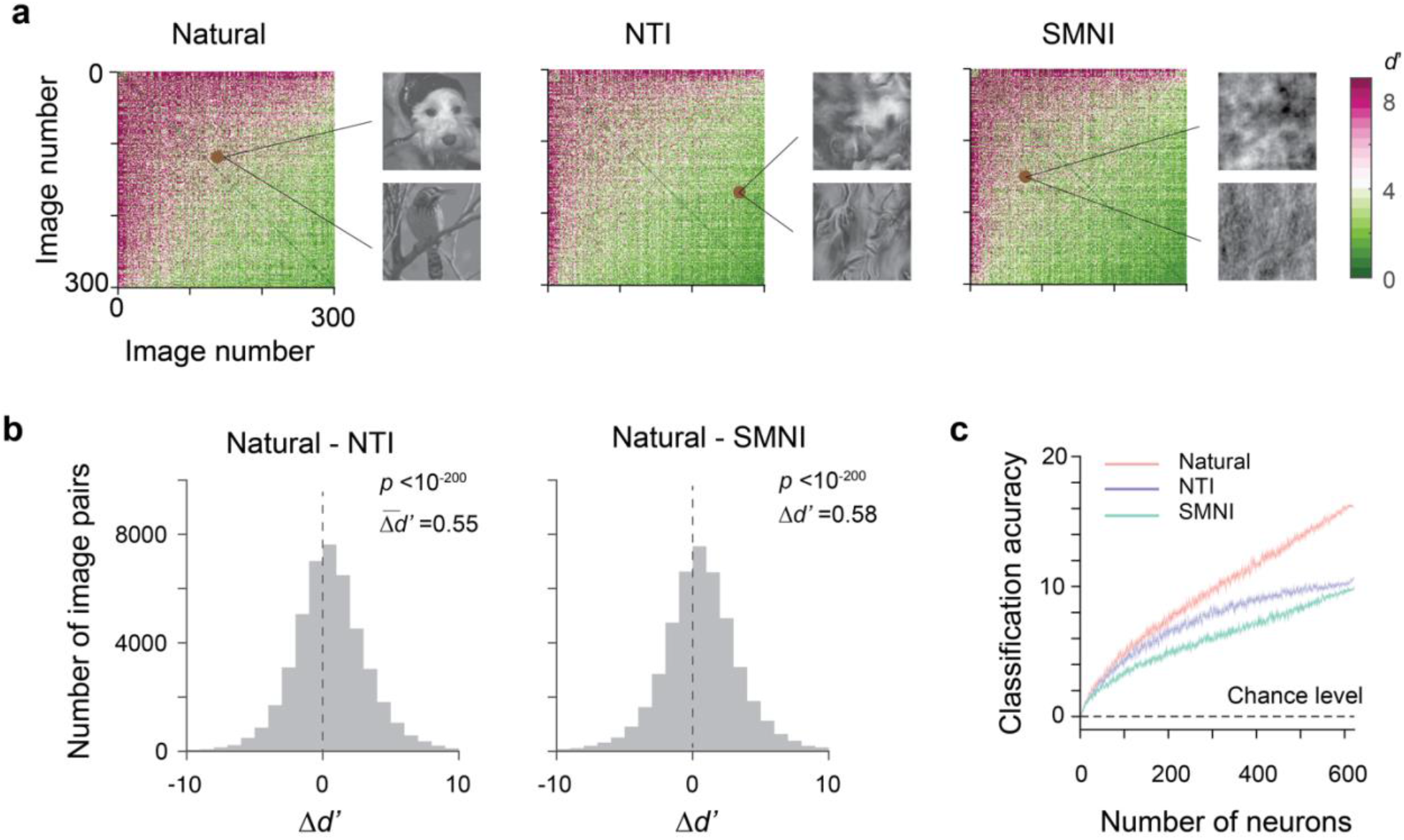
Population representations of different image sets. (**a**). Population discriminability matrix for two pairs of images in different image sets. The brown dots show a representative image pair in the natural image set (left) and its corresponding image pair in the NTI set (middle) and SMNI set (right). (**b**). Population discriminability (d’) for between Natural images and NTIs (left) and between Natural images and SMNIs. (c). Image decoding performance for different image categories as a function of population size. The thickness of lines indicates s.e.m..

We further tested the fidelity of population coding for the different sets of images using image classifiers. We found that a simple nearest-neighbor decoder, trained on half of the trials and tested on the other (Methods), was able to identify the stimulus within the image set with up to 17% classification accuracy, compared to a chance level of 0.3%. We then assessed the decoding performance as a function of the number of neurons. Across different population sizes, classifiers consistently exhibited the highest performance for natural images and lowest for SMNIs (Fig. 6c). In addition, we observed that, for the natural image condition, the decoding accuracy did not saturate at a population size of 600. This is consistent with a population sparse coding strategy in which fewer neurons are driven by a given stimulus in the natural image set than a stimulus in the other image sets. Therefore, performance would further improve with an even larger number of neurons.

## Discussion

We found that single V1 neurons responded more vigorously to natural images than to naturalistic textures and spectrum-matched noise images. This sensitivity to natural images was observed throughout all cortical layers in V1, including neurons in input granular layers 4C alpha and beta. In addition, V1 neurons also showed high lifetime and population sparseness for natural images and population synchrony. Lastly, for population coding, the neuronal population exhibited higher discriminability for natural images. Taken together, our results demonstrate a rapid and pervasive sensitivity to natural images is present at the earliest stages of cortical processing. Below, we discuss the significance and limitations of our observations in three different aspects.

### Relationship to previous studies

Our results integrate two lines of work on natural image representation in V1 in the same experiments. On the one hand, consistent with previous studies on natural texture images ^14,17,18^, our results show that single V1 neurons does not show significant selectivity to naturalistic textures. Specifically, V1 neurons do not show response differences between naturalistic textures and spectrum-matched noise images even with a large population of simultaneously recorded V1 neurons. In addition, the results expand on this previous observation by showing that V1 neurons are not only insensitive to naturalistic textures generated from natural texture images ^14,17,18^, but also to those generated from natural images. However, it would be presumptuous to conclude that V1 neurons are insensitive to any natural image statistics based on their insensitivity to natural textures. Perhaps, in agreement with other studies that compare neuronal responses to natural and SMNIs ^19–23^, our results show that V1 neurons indeed exhibit a robust preference for natural images to SMNIs. In addition, our results reveal a pervasive coding preference for natural images compared to NTIs, both in terms of the involvement of neurons across layers and in terms of the response magnitude. Our experiment integrates two lines of experimental paradigms by testing all three sets of images with different naturalness levels using large scale neuronal recordings in the same animals and by quantifying the neuronal response toward different image sets with the same metrics. Our results support both results and show that V1 neurons are sensitive to the high order image statistics that differ between natural images and NTIs, however, are insensitive the mid-level image statistics that differ between the NTIs and SMNIs.

### Computational benefits of sparseness

A sparse coding strategy refers to the encoding of sensory information using only a small number of active neurons at any point in time ^26^, and it comes in two forms ^27^: lifetime sparseness and population sparseness. Lifetime sparseness describes the activity of a single neuron over time, and population sparseness describes the activity of a population of neurons for a given time window. Previous theoretical, computational, and experimental studies suggested at least four reasons supporting sparse representations as a model of sensory signaling. First, they are useful for storing patterns in memory as they can avoid cross-talk ^38^, and facilitate learning associations in neural networks using local learning rules ^39^. Second, they form neural representations with higher degrees of specificity, making the structure in natural sensory input explicit ^40^. Third, they produce a simple flattened representation of the otherwise curved manifold structure of data, and consequently it is easier to read out at subsequent levels of processing ^41,42^. Fourth, they are energy efficient ^43,44^. A ubiquitous property of primary sensory cortical areas is that they overrepresent their sensory inputs many times over compared to their thalamic input. Experimental evidence for sparse coding has been found in several different sensory cortices, including visual cortex ^21–23,28,30,31^, auditory cortex ^45,46^, olfactory cortex ^47,48^, the primary somatosensory cortex ^49,50^, and others. Theoretical studies suggest that V1 neurons are at least 200 times more abundant than their thalamic input. As a result, V1 neurons could be highly specialized in their feature selectivity and thus highly sparse in their population responses ^51,52^. Consistent with these studies, our results demonstrate for a given number of images, significantly more response variance was explained by natural images than synthetic images, which suggests a higher lifetime sparseness in the coding of natural images by V1 neurons. In addition, for a given number of neurons, significantly more response variance was explained with natural images than with synthetic images, which suggests population sparseness. Moreover, on a finer time scale, V1 neurons were less synchronized with the overall population, which may be consistent with greater sparseness in V1 responses to natural images. Lastly, our results also show that natural images were reliably represented despite the fact that the encoding of natural images utilized a relatively small number of highly active neurons at any point in time.

### Origins of early sensitivity to natural images

We observed a short latency sensitivity to natural images among V1 neurons; sensitivity emerged within 60 ms. The short latency preference for natural images could perhaps arise *de novo* within the earliest stages of V1 processing. Alternatively, V1 neurons might inherit a natural image bias from the retina or dLGN. Previous work demonstrates that retinal ganglion cells are biased toward the coding of contrast decrements, as opposed to increments, reflecting a dark bias in the pixel distribution of natural images ^53^. Similarly, V1 neurons have non-uniformities in their representation of cardinal orientations in ferret and cat ^54,54,55^, which matches a known characteristic of natural images ^56^. We hypothesize that the sensitivity that we observed in V1 may result from the adaptation to the high order statistical features of the natural visual environment. Future research will be needed to address what specific high order statistical features within natural images drive the neuronal coding preference toward natural images in V1.

In addition, previous studies show that natural images can be reconstructed by the population activity of dLGN neurons ^57,58^ and retinal ganglion cells ^59–62^. Specifically, two recent studies suggested that natural image statistics can be used as a prior to improve image estimates in reconstruction methods using retinal neurons ^60,61^, similar to what has been found in visual cortex ^63^. Nevertheless, it is still unknown if neurons in the primate retina or dLGN show stronger response to natural images compared to low- and mid-level image statistics matched synthetic images. Future work will be needed to establish the source of the natural image selectivity we observed in V1.

## Methods

### Subjects

All experimental procedures were in accordance with the National Institutes of Health Guide for the Care and Use of Laboratory Animals, the Society for Neuroscience Guidelines and Policies, and Stanford University Animal Care and Use Committee policies. Anesthetized recordings were conducted in two adult male macaques (13kg and 8kg). The number of animals used is typical for primate neurophysiological experiments. In order to allow for neurophysiological recordings, both monkeys were surgically prepared with head restraint recording chambers under isoflurane anesthesia. Analgesics were used to minimize discomfort. After recovery, anesthetized recordings were subsequently conducted.

### Visual stimulation generation and presentation

We used a large set of images that varied in their higher-order statistics. These images included 300 natural images (NA) (CIFAR-10), 300 naturalistic texture images (NTIs), and 300 spectrum-matched noise images (SMNIs). For each natural image, the corresponding synthetic natural texture image was generated by an iterative optimization procedure that matched the texture structure in the original natural images^15,18^. The spectral matched noise image was generated by randomizing the phase, but matching the two-dimensional (2D) power spectrum. As a result, images in each triplet were identical in their mean luminance, pixel standard deviations, and root mean square (RMS) contrasts (Supplementary Fig. 1).

Visual stimuli were presented on a LCD monitor NEC-4010 (Dimensions= 88.5 (H)* 49.7 (V) cm, pixels=1360 * 768, frame rate= 60 Hz) positioned 114 cm from the monkey. Each image was 2 dva × 2 dva and was presented ten times in a pseudorandom order at the population RF location. In each trial, an image was presented was presented for 0.2s, followed by an 0.25 sec equal luminance blank screen inter-trial interval. Neuronal responses were measured during 9000 trials, with 10 trials per image, during each recording.

### Electrophysiological recordings

Prior to each recording session, treatment with dexamethasone phosphate (2 mg per 24 h) was instituted for 24 hours to reduce cerebral edema. After administration of ketamine HCl (10 mg per kilogram body weight, intramuscularly), monkeys were ventilated with 0.5% isoflurane in a 1:1 mixture of N_2_O and O_2_ to maintain general anesthesia. Electrocardiogram, respiratory rate, body temperature, blood oxygenation, end-tidal CO_2_, urine output and inspired/expired concentrations of anesthetic gases were monitored continuously. Normal saline was given intravenously at a variable rate to maintain adequate urine output. After a cycloplegic agent was administered, the eyes were focused with contact lenses on a CRT monitor. Vecuronium bromide (60 μg/kg/h) was infused to prevent eye movements.

With the anesthetized monkey in the stereotaxic frame, an occipital craniotomy was performed over the opercular surface of V1. The dura was reflected to expose a small (~3 mm2) patch of cortex. Next, a region relatively devoid of large surface vessels was selected for implantation, and the Neuropixels probe was inserted with the aid of a surgical microscope. Given the width of the probe (70 um x 20 um), insertion of it into the cortex sometimes required multiple attempts if it flexed upon contacting the pia. The junction of the probe tip and the pia could be visualized via the (Zeiss) surgical scope and the relaxation of pia dimpling was used to indicate penetration, after which the probe was lowered at least 3-4 mm. Prior to probe insertion, it was dipped in a solution of the DiI derivative FM1-43FX (Molecular Probes, Inc) for subsequent histological visualization of the electrode track.

Each probe consisted of 986 contacts distributed across 10 mm, of which 384 contacts could be simultaneously selected for recording. Probes were inserted into the lateral operculum of V1 at angles nearly perpendicular to the cortical surface under the aid of a surgical microscope. Given the length of the probe (1 cm), and the complete distribution of electrode contacts throughout its length, recordings could be made either in the opercular surface cortex (M1) or within the underlying calcarine sulcus (M2), by selecting a subset of contiguous set of active contacts (n = 384) from the total number (n=986). Of the total, 622 neurons (52%) (M1: 470 neurons, session 1: 280, session 2: 190; M2: 152 neurons, session 3: 83, session 4: 69) were visually responsive throughout the recording session. Receptive fields (RFs) from online multi-unit activity were localized on the display using at least one eye. Recordings were made at 2 sites in one hemisphere of each monkey. At the end of the experiment, monkeys were euthanized with pentobarbital (150 mg kg^−1^) and perfused with normal saline followed by 1 liter of 1% (wt/vol) paraformaldehyde in 0.1 M phosphate buffer, pH 7.4.

### Receptive field mapping

We mapped the location and size of receptive fields (RFs) by pseudorandomly presenting black and white squares within a 10 by 10 probe grid extending 4 by 4 dva. In each recording session, we placed the probe grid to cover the area where we expected most of the RFs, presenting the probe sqaures in random order for 200 ms each. To map the receptive fields of individual neurons, we averaged the activity of a neuron from 30 ms to 150ms after stimulus onset across all repetitions for each location. The receptive fields are defined as the locations where the neuronal response to the stimulus was significantly larger than baseline activity (permutation test, n = 1000, corrected by cluster-based correction for multiple comparisons^64^). Neuronal receptive fields (RFs) spanned eccentricities between ~4 and 6° degree of visual angle (dva) (M1) and ~ 5-7° dva (M2) in the lower visual field; images were positioned on the average RF location.

### Layer assignment

The laminar location of our recording sites was estimated based on a combination of functional analysis and histology results. For each recording, we first performed a current source density (CSD) analysis on the stimulus-triggered average of local field potentials (LFP). LFP signals recorded from each 4 neighboring channels were averaged and realigned to the onset of visual stimulus. CSD was estimated as the second-order derivatives of signals along the probe axis using the common five-point formula ^65^. The result was then smoothed across space (σ = 120 μm) to reduce the artifact caused by varying electrode impedance.

We located the lower boundary of the major sink (the reversal points of sink and source) as the border between layer 4C and layer 5/6. Based on this anchor point, we assigned other laminar compartment borders using the histological estimates. Cortical layers were divided into four comparably sized laminar compartments, specifically layers 2/3, 4A/B, 4C, and 5/6 (Mean depth: 650μm, 311μm, 281 μm, 489μm, respectively). Layer4C was further divided into two 2 input sublaminae. The top half of Layer 4C was assigned as Layer 4C alpha and the bottom half was assigned as Layer 4C beta.

### Image statistics

The room-mean square (rms) contrast is defined as below^66^.

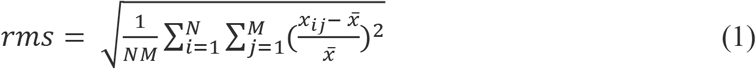

Luminances *x_ij_*([0,255]) refer to the *i*-th *j*-th element of the image of size N by M. 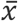 is the average luminance of the image. We also adopted this equation to estimate the local RMS contrast within the sub image of 0.5 dva by 0.5 dva region centered on each pixel of the image^67^.

The band-limited Contrast^66^ was defined in the Fourier domain as:

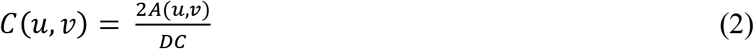

where A(u,v) is the amplitude of the Fourier transform of the image, u and v are the horizontal and vertical spatial frequency coordinates, respectively, and DC is the zero-frequency component.

### Modulation Index

We used modulation indices to quantify the response differences between the image types^14,18^:

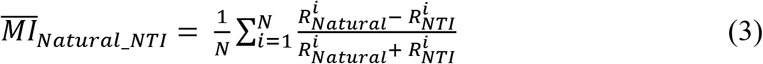

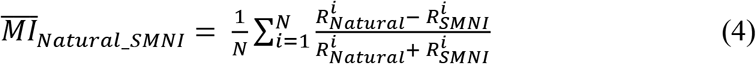

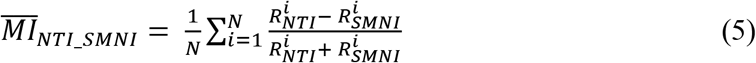

With 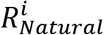 denote the mean neuronal response to *i*th natural image, [40, 200] ms, [40, 100] ms, and [100, 200] ms, relative to stimulus onset for the All, Early and Late periods respectively. N is the total number of images for each image categories.

### GLM encoding models

Gaussian GLM models were used to identify the categorical effect of synthetic texture image and spectrum matched image while controlling the image statistic including local RMS contrast and band-limited contrast. For each image *j*, we divided the local RMS contrast for each image into nine 0.67 dva by 0.67 dva patches and computed the mean RMS contrast for each patch *i rms_i_*, *i* ∈ [1,9]. Similarly, we divided the band-limited contrast into four sections covering 45° and computed the mean contrast for each section i *c_j_*, *j* ∈ [1,4]. The mean response of each neuron in 30 ms to 200ms after image onset was first z-normalized and then modeled as:

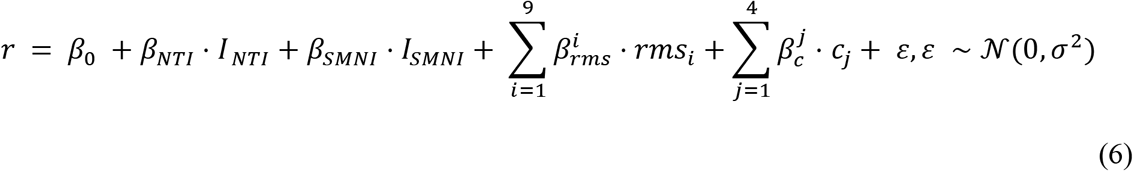

where *I_NTI_* and *I_SMNI_* are dummy variables (0 or 1) for NT images and SMN images respectively, *β_rms_* are regression coefficient for local RMS contrast, *β_c_* are regression coefficient for band-limited contrast, *ε* is a additive noise variable and 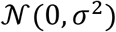 is a Gaussian distribution with mean 0 and variance *σ*^2^. We calculated the time course of GLM coefficient by computing the regression coefficents for each time bins. Spike times were converted to firing rate estimates by convolution with a causal 30 ms boxcar filter with an interval of 5 ms.

We used the scikit-learn 0.24.1 library in Python to fit Poisson GLMs. Poisson GLMs (linear non-linear models) were also used in the analysis. The results are qualitatively similar to Gaussian GLM. We reported the results of the Gaussian GLM in the manuscript, as they are a more robust fit when calculating GLM coefficients across time bins from stimulus onset.

### Measurement of sparseness

We focused on sensory responses by projecting out the dimensions corresponding to baseline activity before further analysis^68^. Specifically, the top 25 dimensions of ongoing activity were found by performing a PCA on the z-scored baseline neural activity recorded during 50ms to 0ms before image presentation. To remove these baseline driven responses, the normalized stimulus driven activity (average neural activity during 40 ms to 200 ms after image representation) was first projected to these top 25 dimensions. This projected activity was then subtracted from the stimulus driven activity.

We calculated the lifetime and population sparseness in two ways. Frist, the sparseness is calculated as the reduced kurtosis of the response distribution, which has previously been used in a number of theoretical and experimental studies^23,40,69^

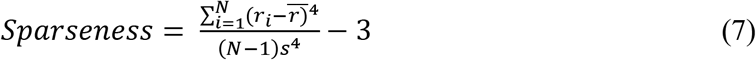

For lifetime sparseness, *r_i_* is the response of a neuron to *i*th image within an image category, 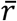 is the mean response of a neuron to all image stimulus within an image category, and N is the number of images for each image category. For population sparseness, *r_i_* is the response of *i*th neuron to a single image stimulus within an image category, 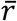 is the mean response across the population of neurons to an image, and N is the number of neurons recorded. Population sparseness was then averaged across images and lifetime sparseness was averaged across neurons.

Second, to examine the sparseness using the whole distribution of the date. We calculated the response *r_i,j_* of each neuron *i* to each image *j*. For lifetime sparseness, we sorted the responses variance explained by each image from high to low. We then measured the sparseness by computing the cumulative variance as a function of number of images. We expected to observe higher variance explained by fewer images if the lifetime sparseness is high. To compare the number of images is needed to capture 80% of the total variance, we randomly sampled 250 images out of 300 images for 500 time. Each time we calculated the number of images are needed capture 80% of the total variance and compared between different imaged categories. Similarly, for population sparseness, we sorted the response variance explained by each neuron from high to low.

The sparseness is quantified as a function of number of neurons for population sparseness. We expected to observe higher variance explained by fewer neurons if the population sparseness is high. To compare the number of neurons needed to capture 80% of the total variance, we randomly sampled 500 neurons (out of 622) for 500 iterations. Each time we calculated the number of neurons needed capture 80% of the total variance and compared between different imaged categories.

### Cross correlation analysis

We computed cross-correlations of spiking activity between each single neuron’s activity and population neuronal activity recorded simultaneously for each experimental session. Correlations were computed using a jitter correction method, which corrects for slow temporal correlations and for stimulus-locked correlations (44, 45). The jitter-corrected correlations were computed by subtracting the expected value of correlations produced from a resampled version of original spike trains, with spike-times randomly shuffled (jittered) within a specified temporal window (the jitter window). The correction term is the average over all possible resamples of the original spike trains, and is subtracted from the raw correlation. Empirical correlations were computed using small discrete time bins Δt, and spikes were jittered within larger jitter bins T. The jittering technique preserves two marginals: the total spike count in each time bin Δt summed across all trials (the peristimulus time histogram, PSTH), and the spike count within each jitter bin T on each trial (the instantaneous firing rate computed in bins T). For each spike on each trial, a new spike is chosen randomly from the set of all spikes within the same jitter bin on all of the trials. Jitter correction removes correlations on timescales greater than the jitter window T. Because it preserves the PSTH shape, jitter correction also removes correlations due to stimulus-locked firing rate modulation.

We computed cross-correlation for stimulus driven activity during 0 to 200ms after stimulus onset. Correlations was computed with Δ*t* = 5*ms*, using a jitter window of T = 50ms. For each neuron, the strength of the population coupling is defined as the peak magnitude of the cross-correlation and the time lag of the population coupling is defined as time lag of the peak.

### Computation of population *d*′ for neural responses to visual stimuli and Image decoder

To estimate how much information the neural activity conveyed about the stimulus identity, we used the population *d*′, which characterizes how readily the distributions of the neural responses to the two different sensory stimuli can be distinguished ^35^. We computed the population *d’* for each image pair within the 300 images for each image categories. A challenge was that calculation of population *d’* in a large population vector space would have involved the estimation of an *N* × *N* noise covariance matrix ^36^. Direct estimation of the covariance matrix would have been unreliable, because the number of neurons (622), was much larger than the number of trial (n=10). We therefore used PCA dimensionality reduction using the trail averaged data for each image within an image set to find the dimensions of the population vector space that preserve the data’s variation across different images. We projected the high dimensional ensemble neural response (N= 622) to a truncated set of dimensions *N_r_* ∈ [1,10] identified by the PCA analysis and calculated the population *d’* for each image pairs.

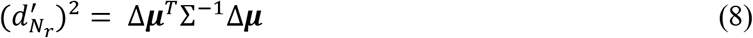

Where 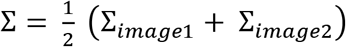 is the noise covariance matrix average across two image stimulation conditions, Δ***μ*** = ***μ***_*image*1_ – ***μ***_*image*2_ is the vector difference between the mean ensemble neuronal responses to the two images.

To decode the stimulus identity from the neural responses, we built a simple nearest-neighbor decoder based on correlation ^68^. We used the average response of the odd trials as the training set while using the average response of the even trials as the test set. We correlated the population responses for an individual stimulus in the test set with the population responses from all stimuli in the training set. The stimulus with the maximum correlation was then assigned as our prediction. We defined the decoding accuracy as the fraction of correctly labelled stimuli.

## Acknowledgments

We thank Tim Harris and Karel Svoboda for providing the Neuropixel probes, Jonathan C. Horton for extensive help with the recordings and histology, E.J. Chichilnisky and Kenneth H. Britten for helpful comments on the results and interpretations, Wenqing Hu for help with the data analysis, and Shellie Hyde and Sam Baker for technical assistance. This work was supported by NIH Grant EY014924 and EY029759.

## Author contributions

X. C., A. M. N., and T. M. designed the research. X. C., S. Z., and T. M. performed the experiments. X. C., S. Z., K. B., R. X., N. K., A. M. H., and T. M. analyzed the data. X.C., and T. M. wrote the paper.

---

**Figure S1.**
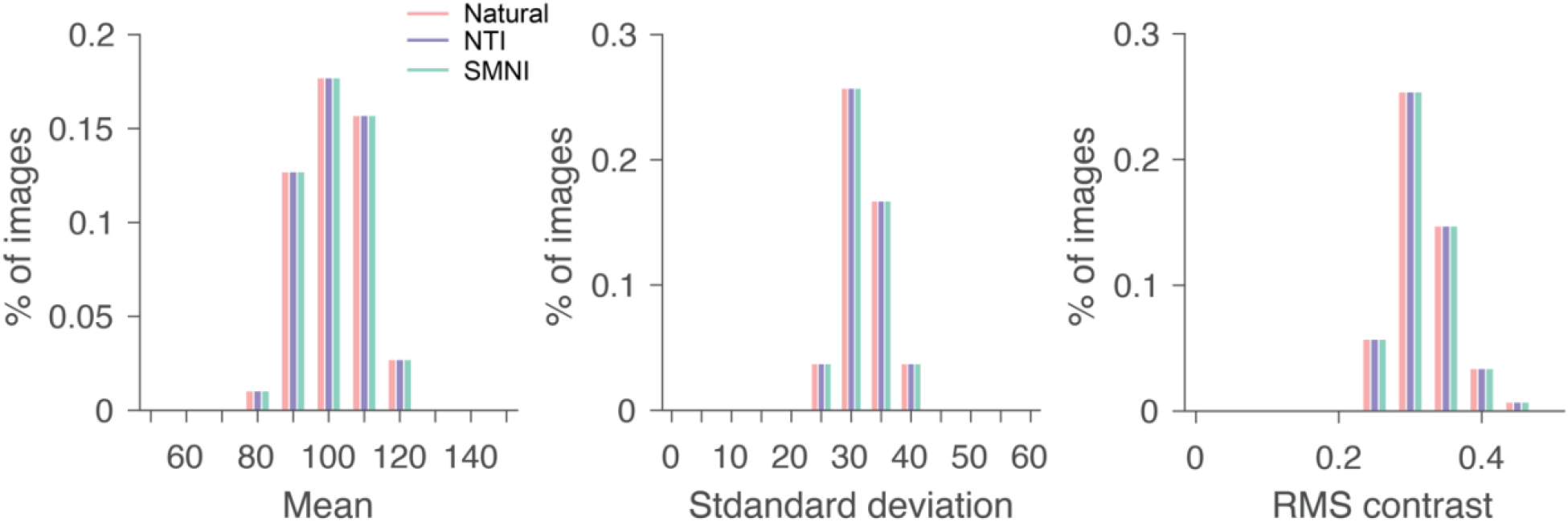
Comparison of global image statistics across different image sets. Left, mean luminance; Middle, pixel standard deviations; and Right, RMS contrast of three image sets.

**Figure S2.**
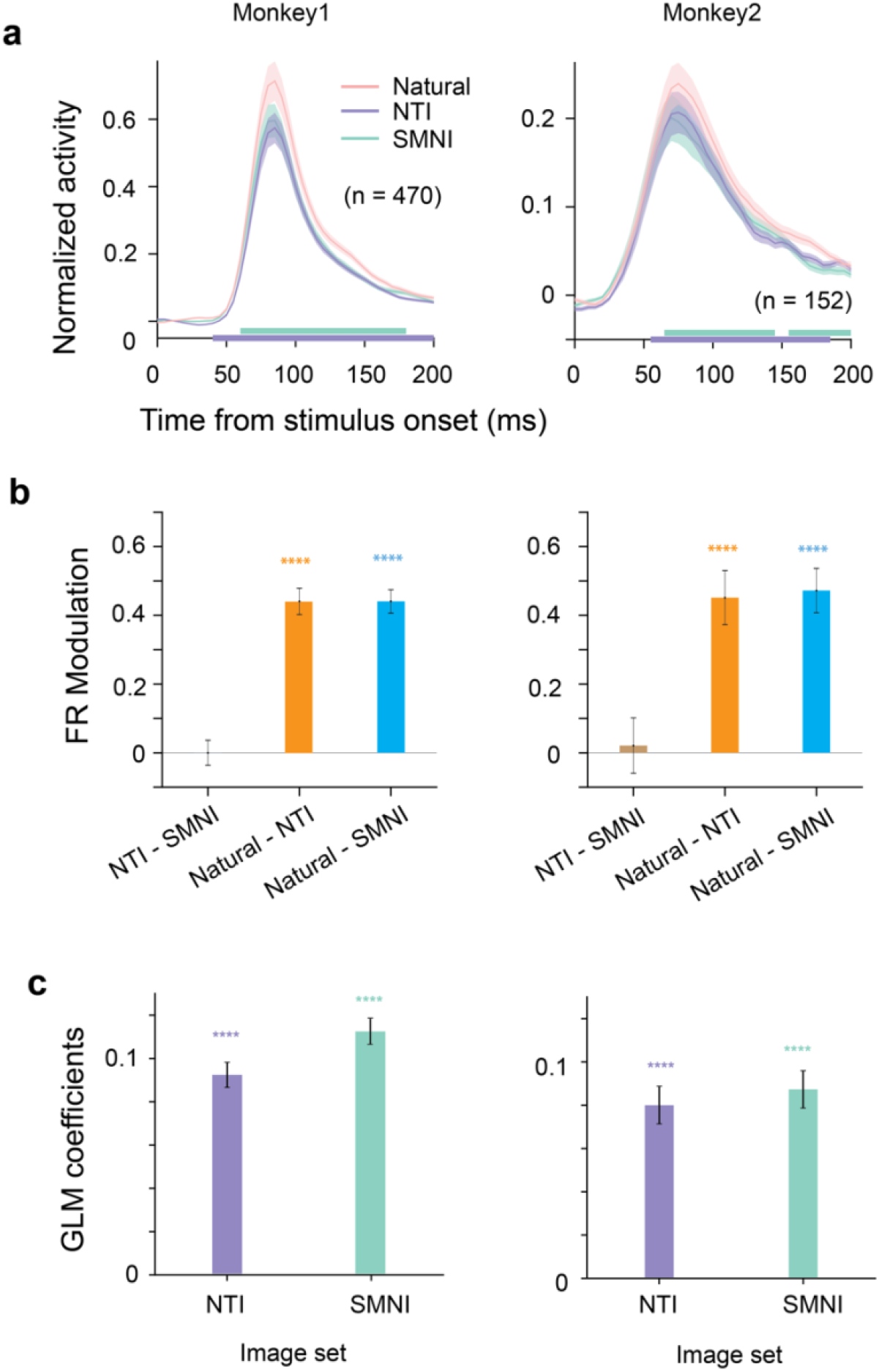
Neuronal responses to different image categories for both monkeys. (Monkey 1: Left, Monkey 2: Right) (**a**) Time course of average normalized firing rate across all recorded neurons in V1 to images of natural image (pink), NTI (purple), and SMNI (green). The same conventions were used as in **Fig. 2a**. (**b**) Box plots summarize the modulation index for comparison between different image sets using the whole response epoch (all: 40 – 200ms). (**c**) Box plot summarizes the GLM coefficients for synthetic texture images (purple) and spectrally matched noise (green). **** denotes *p* < 10^−4^.

**Figure S3.**
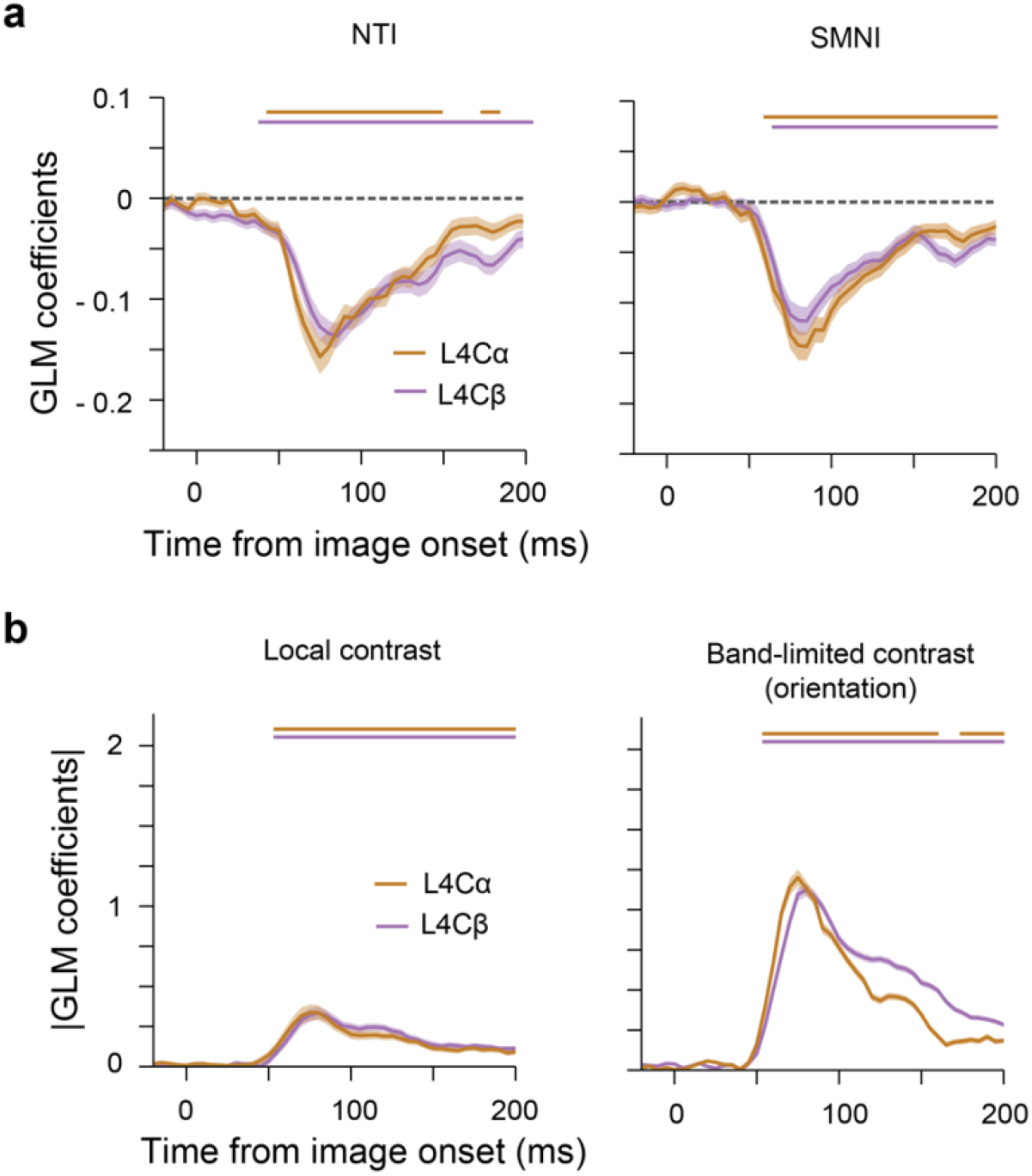
Comparison between sublaminae L4C*α* and L4C*β*. (a) Average GLM coefficients for NTI images (left) and SMNI images (right) as a function of time, separated by different L4C sublaminae. The thickness of lines indicates the s.e.m. of GLM coefficients across neurons. The lines above indicate the time periods when the coefficients are significantly different from zero. (b) Average GLM coefficients for local contrast (left) and orientation (right) as a function of time, separated by different sublaminae. Horizontal lines indicate the time periods when the coefficients are significantly different from zero (p < 0.05, corrected for multiple comparisons.

**Figure S4.**
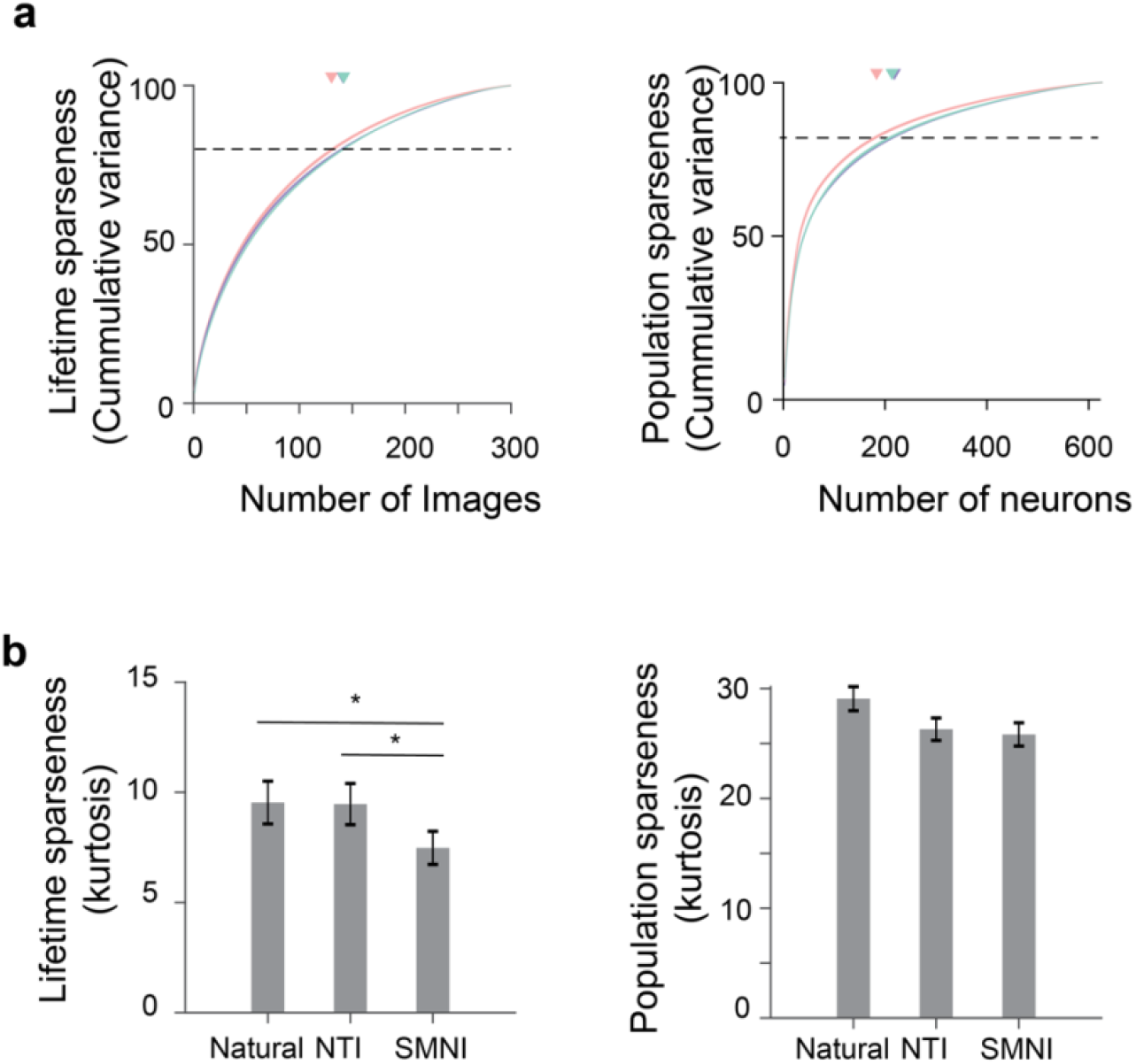
Lifetime and population sparseness. Lifetime and population sparseness was quantified in two ways: (a) Cumulative variance as a function of number images (left: Lifetime sparseness) and number of neurons (Right: population sparseness). (b) Reduced kurtosis of the response distribution (left: Lifetime sparseness; Right: Population sparseness). * denotes *p* < 0.05 (paired t-test).

**Figure S5.**
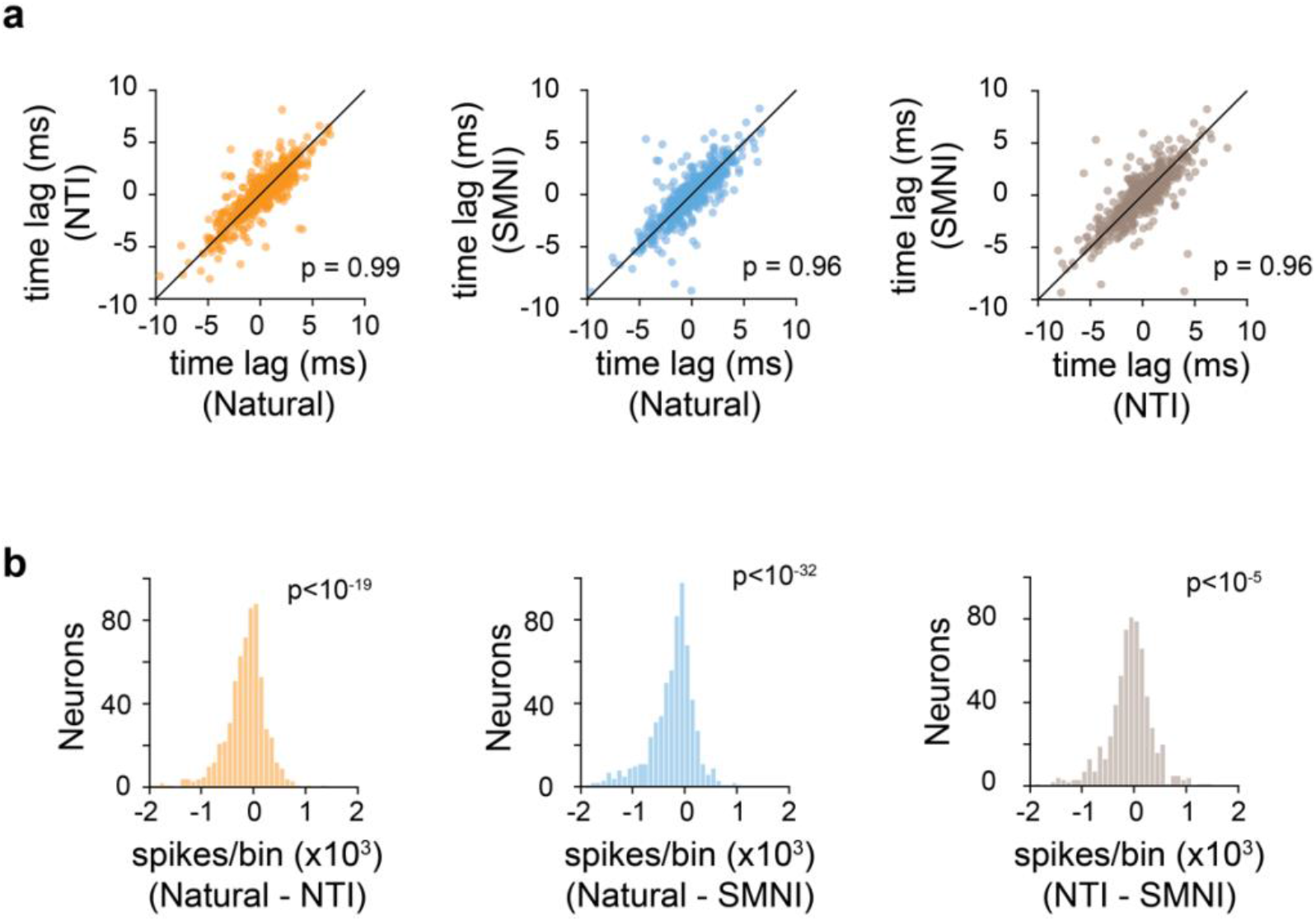
Population synchrony for different image categories. (**a**) Scatterplots compare the time lags of the population coupling. Left, comparison between natural images and NTIs; Middle, comparison between natural images and SMNIs; Right, comparison between NTIs and SMNIs. (**b**) Histograms compare peak magnitude of the population coupling. Left, comparison between natural images and NTIs; Middle, comparison between natural images and SMNIs; Right, comparison between NTIs and SMNIs. *p* values were computed using paired t-tests.

**Figure S6.**
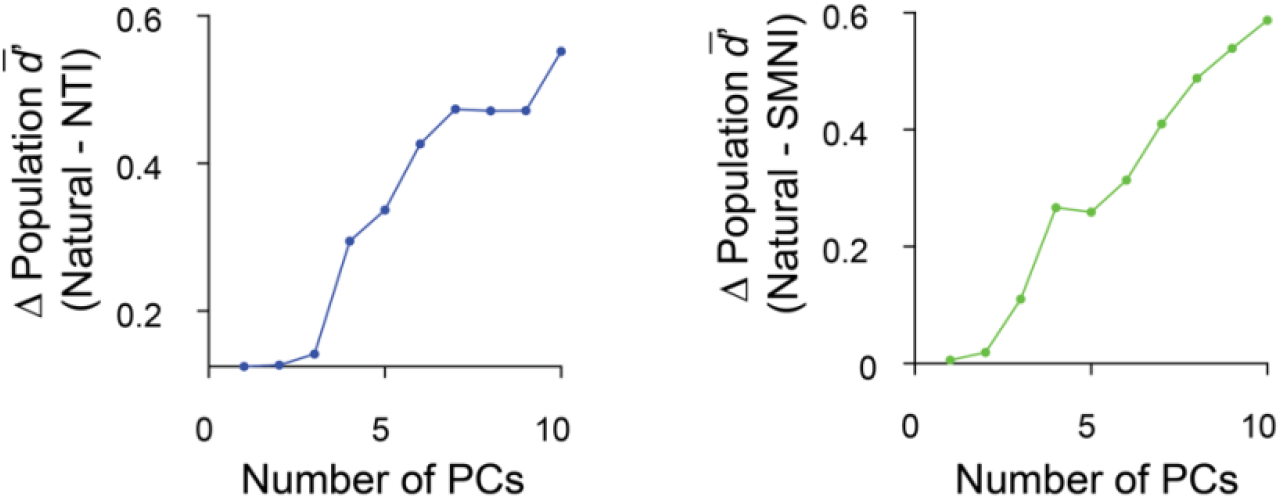
Differences of population *d*′ as a function of PC dimensions. Left, differences of population *d’* between natural images and NTIs; Right, differences of population *d’* between natural images and SMNIs.

